# The VarA-CsrA regulatory pathway influences cell shape in *Vibrio cholerae*

**DOI:** 10.1101/2021.09.09.459595

**Authors:** Leonardo F. Lemos Rocha, Katharina Peters, Jamie S. Depelteau, Ariane Briegel, Waldemar Vollmer, Melanie Blokesch

**Author notes:** **Author Contributions:** L.F.L.R and M.B. designed the research; L.F.L.R. performed most of the experiments; L.F.L.R. and M.B. analyzed the data; K.P. and W.V. performed the peptidoglycan composition analysis; J.S.D. and A.B. performed the cryoelectron microscopy; L.F.L.R and M.B. wrote the manuscript with input from K.P., J.S.D., A.B., and W.V. All authors approved the manuscript. **Competing Interest Statement:** The authors declare no conflict of interest.

## Abstract

Despite extensive studies on the curve-shaped bacterium *Vibrio cholerae*, the causative agent of the diarrheal disease cholera, its virulence-associated regulatory two-component signal transduction system VarS/VarA is not well understood. This pathway, which mainly signals through the downstream protein CsrA, is highly conserved among gamma-proteobacteria, indicating there is likely a broader function of this system beyond virulence regulation. In this study, we investigated the VarA-CsrA signaling pathway and discovered a previously unrecognized link to the shape of the bacterium. We observed that *varA*-deficient *V. cholerae* cells showed an abnormal spherical morphology during late-stage growth. Through peptidoglycan (PG) composition analyses, we discovered that these mutant bacteria contained an increased content of disaccharide dipeptides and reduced peptide crosslinks, consistent with the atypical cellular shape. The spherical shape correlated with the CsrA-dependent overproduction of aspartate ammonia lyase (AspA) in *varA* mutant cells, which likely depleted the cellular aspartate pool; therefore, the synthesis of the PG precursor amino acid meso-diaminopimelic acid was impaired. Importantly, this phenotype, and the overall cell rounding, could be prevented by means of cell wall recycling. Collectively, our data provide new insights into how *V. cholerae* use the VarA-CsrA signaling system to adjust its morphology upon unidentified external cues in its environment.

**Significance Statement:** Responsible for the diarrheal disease cholera, the bacterium *Vibrio cholerae* tightly regulates its virulence program according to external stimuli. Here, we discovered that a sensing-response mechanism involved in the regulation of virulence also controls bacterial shape. We show that *V. cholerae* lacking this system lose their normal comma shape and become spherical due to an abnormal cell wall composition caused by metabolic changes that reduce available cell wall building blocks. Our study therefore sheds new light on how *V. cholerae* modulates its morphology based on environmental changes.

## Introduction

The current ongoing 7^th^ cholera pandemic sickens millions of people every year (1), though many questions still surround the pathogenicity of its causative agent, the well-studied gram-negative bacterium *Vibrio cholerae. V. cholerae* is frequently found in aquatic habitats (2), but throughout human history, it has caused several cholera pandemics, leading to questions about how it switched from an environmental to a pathogenic lifestyle. Nonetheless, it is well established that virulence induction is linked to the ability of the bacterium to sense its surroundings (3). Hence, it is important to study the mechanisms that allow bacteria to detect environmental changes and rapidly adapt to them.

One example of such a sensing mechanism is the **v**irulence-**a**ssociated **r**egulators S and A (VarS/VarA) two-component system (TCS) of *V. cholerae*. For this TCS, the sensor kinase VarS detects an unknown signal and activates the response regulator VarA through phosphotransfer. Subsequently, phosphorylated VarA binds to the promoter regions of the genes that encode the small RNAs (sRNAs) *csrB, csrC*, and *csrD*, fostering their transcription. These sRNAs control the activity of carbon storage regulator A (CsrA) by sequestering it away from its mRNA targets (4). This sRNA-based sequestration mechanism is highly conserved among gamma-proteobacteria and known to be important beyond virulence regulation in those organisms (5). CsrA is a post-transcriptional regulator that binds to consensus motifs of specific mRNAs, thereby controlling access to the ribosome binding site, which ultimately promotes or prevents translation (6). Additionally, CsrA controls the formation of RNA hairpins, which can expose Rho-binding sites, leading to premature transcriptional termination. Furthermore, CsrA controls mRNA stability by preventing mRNA cleavage by the endonuclease RNase E (6) and it has the flexibility to bind mRNAs in different conformational states, thereby increasing the number of genes that can be modulated by this global regulator (7).

RNA-based regulatory systems often control virulence in pathogenic gamma-proteobacteria (8). Indeed, for *V. cholerae*, the VarS/VarA system was regarded as a virulence-associated regulatory TCS, given that a *varA* mutant produced reduced levels of the two major virulence factors (e.g., the cholera toxin and the toxin co-regulated pilus) compared to its parental wildtype (WT) strain. This reduced production of virulence factors resulted in an *in vivo* fitness disadvantage of the mutant compared to the WT upon intestinal colonization of infant mice (9). The VarS/VarA system also contributes to the dissemination of *V. cholerae* from the host into the environment (10). Moreover, in conjunction with CsrA, this TCS is known to be involved in the regulation of central carbon metabolism, iron uptake, lipid metabolism, flagellum-dependent motility, and other phenotypes (11, 12). Collectively, signaling through the VarS/VarA-CsrA circuit therefore affects the environmental lifestyle of *V. cholerae* as well as its pathogenesis.

The VarS/VarA system is also involved in quorum sensing (QS) (4, 13), which is a cell-to-cell communication process mediated by the secretion, accumulation, and sensing of extracellular signaling molecules (autoinducers). This process fosters synchronized bacterial behavior, such as bioluminescence, biofilm production, competence for DNA uptake, and virulence regulation (14). In *V. cholerae*, the master regulator of QS, HapR, is produced at high cell densities. Previous work has shown that VarA controls HapR abundance, as this QS regulator was undetectable in the absence of *varA* (4, 13). However, the exact mechanism behind this regulation is still not fully understood.

Here, we aimed to better understand the VarA-CsrA signaling pathway to explore the role of this sensing system on the emergence of virulence in *V. cholerae*. In this work, we identified a new role of this regulatory system in the modulation of the cellular morphology of *V. cholerae*. More precisely, we show that its normal morphology is lost in *varA*-deficient strains, resulting in round instead of curve-shaped cells during prolonged growth. This spherical morphology is likely the result of a weakened peptidoglycan (PG) cell wall due to reduced numbers of peptide cross-linkages. Moreover, we demonstrate that the changed PG composition is a consequence of a lack of cell wall precursors due to the overproduction of the aspartate ammonia lyase enzyme. Collectively, this study deciphers how the VarA-CsrA pathway regulates cell shape by modulating cell metabolism, and therefore, cell wall building blocks, giving more insight to the bacterial regulation of a pathway required for virulence.

## Results and Discussion

### VarA deficiency results in cell rounding

Given that VarS/VarA are a conserved regulatory pathway among gamma-proteobacteria (5), we sought to study the contribution of this TCS towards diverse *V. cholerae* cellular processes. Unexpectedly, while generating mutants of this TCS, we observed that *varA*-deficient *V. cholerae* cells exhibited an unusual cell morphology (Fig. 1A). Instead of the comma-like shape common for most *Vibrio* species and that was observed for the parental WT strain, the *varA* mutant cells were found to be round in shape after overnight growth. Complementation of the mutant through the provision of the promoter-preceded *varA* gene *in cis* (ΔvarA+*varA*) restored the WT morphology (Fig. 1A), supporting the causality between VarA deficiency and the changed cell shape.

**Figure 1.**
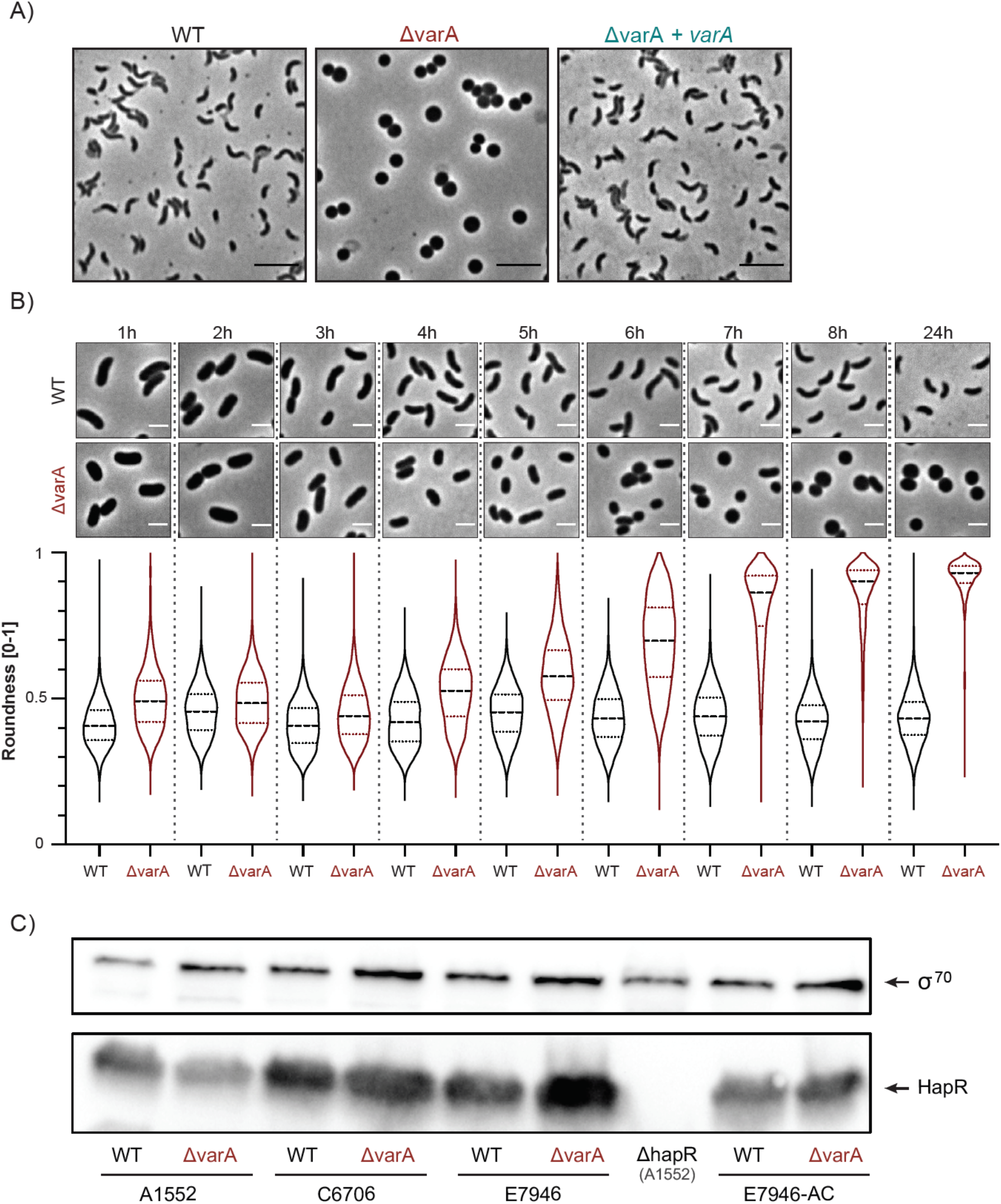
*V*. *cholerae* ΔvarA cells change their morphology at later stages of growth. (A) ΔvarA cells grown overnight are round. Phase contrast micrographs of the WT, ΔvarA, and the complemented ΔvarA+*varA* strains that were grown for 20 h. Scale bar: 5 μm. (B) ΔvarA cells become round during late growth. Phase contrast microscopy imaging (top) and quantification (roundness values at the bottom) of the WT and ΔvarA strains during growth. Cells were imaged every hour for 8 h and again at 24 h post-dilution. Scale bar: 2 μm. The roundness was quantified using the MicrobeJ software and is based on 3,000 cells each (n = 1000 per biologically independent experiment). (C) ΔvarA cells produce the QS regulatory protein HapR. Western blot analysis to detect the HapR protein levels in the different *V*. *cholerae* El Tor pandemic strains A1552, C6707, E7946, and their *varA-*deficient mutants, with the A1552ΔhapR strain as a negative control. The WT and ΔvarA variant of strain E7946 shown on the right were provided by Andrew Camilli (E7946-AC). All strains were sampled at an OD_600_ of ~2.5. Detection of σ^70^ served as a loading control.

To investigate the shape of a larger number of bacteria while reducing image selection bias, we implemented MicrobeJ as a tool for quantitative single cell microscopy image analysis (15). This software can measure the *roundness* of a cell, with a parameter value of 0 representing a straight line and 1 a perfect sphere. The analysis of 3000 WT *V. cholerae* cells showed that the majority of the cells depicted a roundness value of ~0.4 (Fig. S1). In contrast, the round-shaped ΔvarA cells showed values close to 1, confirming a nearly spherical cell shape of the mutant. Upon complementation, the measured parameter decreased towards the values of the WT upon complementation (Fig. S1).

Given that VarA is thought to primarily act downstream of the histidine kinase VarS, we next assessed the morphology of *varS*-deficient cells. Interestingly, this mutant strain did not phenocopy the ΔvarA morphology (Fig. S1), even though an intermediate phenotype was occasionally observed. Additionally, a ΔvarSΔvarA double mutant was morphologically indistinguishable from the *varA* single mutant (Fig. S1). The difference between the phenotypes of the *varA* and *varS* mutants suggests that VarA can act independently of VarS. This finding is consistent with previous work by Lenz *et al*. who demonstrated a 10-fold stronger regulatory effect of VarA compared to VarS and a partially VarS-independent but VarA-dependent regulation of the sRNA genes *csrB* and *csrD* (4). Moreover, it is known that the homologous system GacS/GacA in *Pseudomonas aeruginosa* forms a multicomponent signal transduction system with other histidine kinases (16). As such, additional histidine kinases might also exist in *V. cholerae* that bypass VarS.

### VarA-dependent cell morphology is growth-phase specific

For further insight into the underlying dynamics of the morphological phenotype of the ΔvarA mutant, we examined whether rounding occurred at a particular point during growth by performing time-course experiments. Here, we assessed cellular morphology every hour, including a 24 h control sample. As shown in Fig. 1B, the *varA-*deficient cells initially maintained a WT-like morphology, which started to change around 4 h post-dilution. From this point onwards, the roundness of the cells continuously increased until they reached peak roundness at 8 h. We did observe a growth defect for the ΔvarA strain compared to WT (as shown by the enumeration of colony-forming units [CFU] per ml and OD_600_ measurements; Fig. S2). However, as the WT bacteria maintained their *Vibrio* shape throughout the duration of the experiment, the growth delay could be excluded as the sole reason for the changed morphology of the *varA-*deficient cells (Fig. 1B). Hence, we concluded that the abnormal cell shape of the *varA* mutant was growth-phase dependent and most prominent at later time points during growth. Collectively, these data suggest a previously unknown role of VarA in cell shape maintenance during the stationary phase of *V. cholerae*.

### Quorum sensing (QS) is maintained in *varA*-deficient cells

Given the timing of the observed shape transition phenotype and the reported role of VarA in the production of HapR (4, 13), we asked whether the cell rounding was linked to a HapR deficiency, and therefore, a QS defect. However, to our surprise and in contrast to previous reports (4, 13), the ΔvarA cells still produced copious amounts of the HapR protein (Fig. 1C). We confirmed this finding by deleting *varA* in two additional pandemic O1 El Tor *V. cholerae* strains (C6706 and E7946) and by testing a previously published *varA* mutant from the Camilli laboratory (10) (E7946-AC; Fig. 1C). In addition to the production of HapR in these strains, we observed that deleting *hapR* in *varA*-deficient *V. cholerae* partially reduced the cell rounding phenotype. These results further support the notion that HapR is present and active in ΔvarA cells (Fig. S3A). Notwithstanding, deleting *luxO*, whose gene product indirectly acts as a repressor of HapR synthesis at low cell density, did not change the rounding phenotype of the *varA*-deficient cells (Fig. S3B).

To follow up on the inconsistency of the observed HapR production in *varA*-deficient cells compared to previous studies (4, 13), we wondered if this could be linked to a frequently used QS-impaired lab domesticated variant of *V. cholerae* (strain C6706) (17). In previous work, we found that such QS-impaired domesticated variants differ from wild bacteria, and that much of the research done on these strains does not accurately reflect the behavior of QS-proficient strains (17). Indeed, in the case of the C6706 variant, a gain-of-function (GOF) mutation in *luxO* (encoding LuxO[G333S]) lowers *hapR* transcript levels even at a high cell density, which is known to mask many important phenotypes (17). We therefore introduced the *luxO* GOF mutation into strains A1552 and scored HapR levels in the presence or absence of *varA* (Fig. S3C). For comparison, we also included the original C6706 strain (before it was rendered streptomycin resistant; kind gift from J.J. Mekalanos) and a representative sample of its lab-domesticated variant (named here C6706-mut) and their ΔvarA mutants in this analysis (Fig. S3C). These data showed that the already low HapR levels in those strains that carried the *luxO*[G333S] GOF mutation were indeed diminished beyond the level of detection upon deletion of *varA*, which likely explains the discrepancy between our results and previous reports (4, 13). However, in the presence of native, non-mutated *luxO*, deleting *varA* did not abrogate HapR production. We therefore conclude that a defective QS circuit is unlikely to be responsible for the ΔvarA cell morphology.

### VarA deficiency does not phenocopy the VBNC state

Since the abnormal morphology of *varA*-deficient cells occurred at the stationary phase, we wondered whether cells were entering the viable but non-culturable (VBNC) state. This state is usually triggered by restrained metabolic activity caused by, for instance, low nutrients availability and/or an extended time at low temperatures (18). Cells that enter this state are round in shape due to multiple morphological changes, such as dehiscence of the inner and outer membranes and the resulting enlargement of the periplasmic space (19). To test whether *Δ*varA cells had similar morphological defects, we imaged *varA*-deficient cells and their parental WT cells after 20 h of growth by cryo-electron microscopy. As shown in Fig. S4, neither the size of the periplasm nor the morphology of the membranes appeared significantly altered in the *varA* mutant compared to the WT. Collectively, and taken together with the fact that the bacteria maintained their culturability, these observations suggest that the rounding phenotype of ΔvarA cells differed from the well-described *V. cholerae* VBNC state.

### Spherical-shaped *varA*-deficient cells have an abnormal PG

Given the absence of any obvious morphological defects of the inner and outer membranes and the periplasm, we speculated that the absence of VarA might impair the PG fine-structure, as this mesh-like polymer is known to be required to maintain cell shape (20). The PG is located between the inner and outer membranes in gram-negative bacteria (Fig. 2A) and composed of glycan chains of alternating N-acetylglucosamine (GlcNAc) and N-acetylmuramic acid (MurNAc) residues (Fig. 2A). The poly-GlcNAc-MurNAc glycan chains are connected by peptides and both together form the sacculus around the inner membrane (20). Hence, we isolated the PG from the WT, ΔvarA, and complemented ΔvarA+*varA* strains, digested it with the muramidase cellosyl and analyzed the composition of the muropeptides by High Performance Liquid Chromatography (HPLC). We collected data at two different time-points: (1) at 2 h post-dilution, when the cell shape of the *varA* mutant resembles that of the WT; and (2) at 20 h post-dilution, when the mutant exhibits its round morphology. Interestingly, the analysis of the isolated PG showed an unusual composition in *varA*-deficient cells at 20 h post-dilution (Fig. 2B-D; Table S1). Specifically, a new peak (number 4; red arrow in Fig. 2B) appeared exclusively in the chromatogram of the ΔvarA samples and was strongly increased in the 20 h sample (to 24.4±0.0% of total muropeptides), while this peak was below detection limit in the samples from the wild-type or complemented mutant strain (Fig. 2C, Table S1). Mass spectrometry analysis confirmed this peak as disaccharide-L-Ala-D-Glu (Di) (Fig. 2B, Table S1). The high abundance of these dipeptides (24.4±0.0%) came at the expense of the tetrapeptides, which were significantly reduced in the ΔvarA cells (Fig. 2C and Table S1; from 92.4±5.9% abundance in late-grown WT down to 59.1±3.9% in the late-growth ΔvarA mutant). Importantly, the PG of the 20 h ΔvarA sample had significantly reduced peptide cross-linkage with 34.9±1.1% peptides present in cross-links compared to 51.8±1.7% and 51.9±0.5% in the wild-type and *varA-*complemented strain, respectively (Fig. 2D, Table S1). Hence, we conclude that the round shape of the ΔvarA strain at the late growth stages is the result of a weakened PG caused by the high abundance of dipeptides and reduced peptide cross-linkage (Fig. 2E).

**Figure 2.**
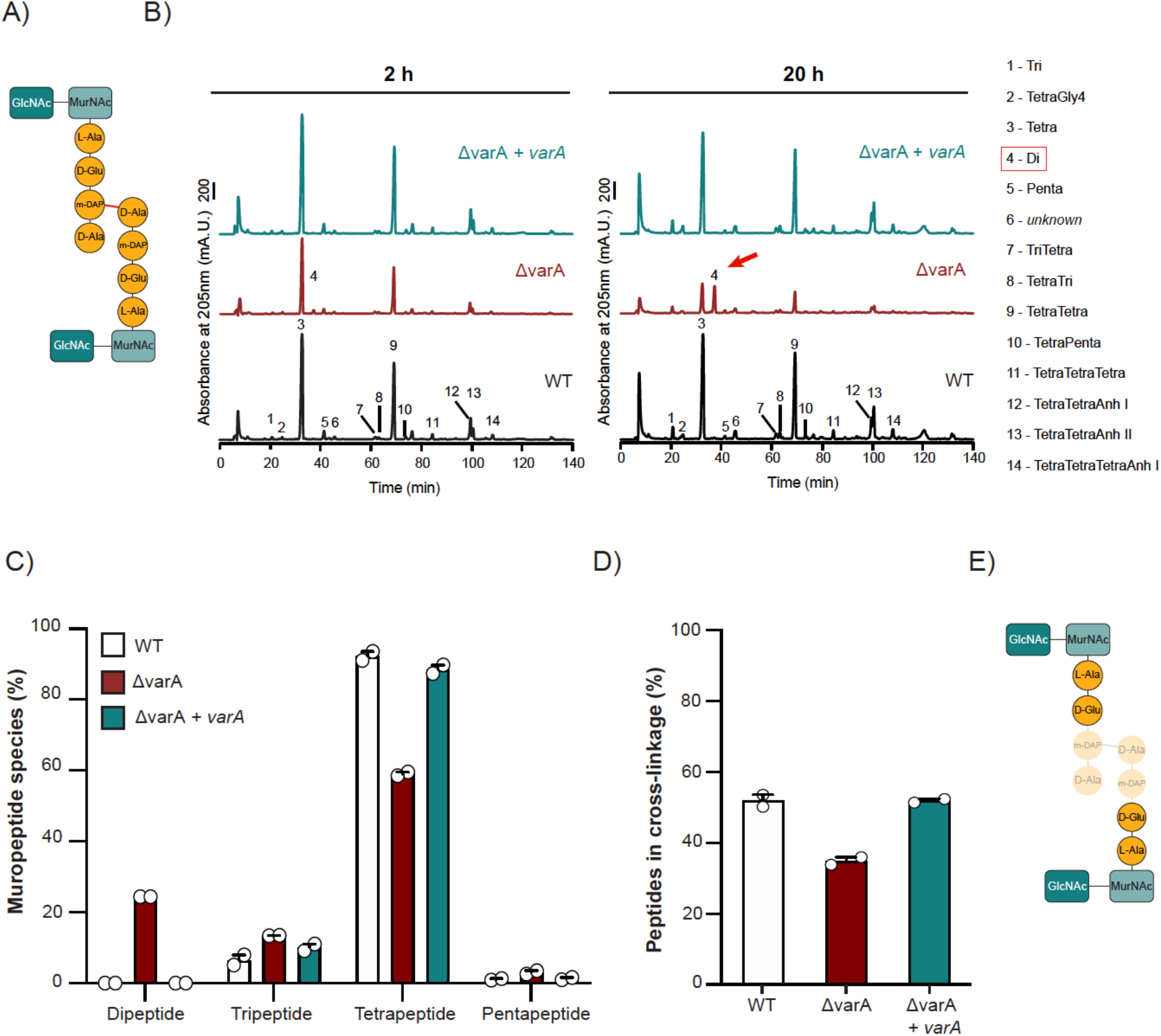
The ΔvarA mutant has an unusual PG composition. (A) Scheme of the peptide cross-link in PG. GlcNAc – N-acetylglucosamine; MurNAc – N-acetylmuramic acid; L-Ala – L-alanine; D-Glu – D-glutamic acid; m-DAP – meso-diaminopimelic acid; D-Ala – D-alanine. (B) The ΔvarA strain contains a new muropeptide and less peptide cross-links. High performance liquid chromatograms of purified PG from the WT, ΔvarA, and complemented ΔvarA+*varA* strains isolated after 2 h or 20 h of growth, followed by digest with cellosyl. Numbered peaks correspond to each muropeptide indicated on the right. (C) The ΔvarA strain contains dipeptides in the PG. Quantification of the different peptides in the purified PG of the WT, ΔvarA, and complemented ΔvarA+*varA* strains isolated at 20 h post-dilution. (D) The ΔvarA strain has less peptide cross-linking. Percentage of cross-linked peptides in purified PG derived from the WT, ΔvarA, and ΔvarA+*varA* strains after 20 h of growth. For (C) and (D): Values are mean (± variance of two biological replicates). See Table S1 for details. (E) Scheme illustrating that dipeptides replace peptide cross-links in the PG of the ΔvarA strain.

To our knowledge, such a high abundance of dipeptides was never observed in *V. cholerae*. One hypothesis to explain this phenotype could be the misregulation of an endopeptidase that cleaves between the second and third amino acids in the peptides. It is known, for instance, that a DL-carboxypeptidase (Pgp1) cleaves disaccharide tripeptides into dipeptides in *Campylobacter jejuni* (21). In *C. jejuni*, the deletion or overexpression of Pgp1 prevents the normal helical morphology of the bacterium as a result of the higher and lower abundance of dipeptides, respectively (21). More recently, Pgp1 was also shown to be involved in the *C. jejuni* helical-coccoid transition after extended incubation or starvation (22). We therefore inspected the annotated genome of *V. cholerae*, but could not identify any obvious DL-carboxypeptidase gene. Interestingly, the amidase AmiA of *Helicobacter pylori* was also shown to foster an accumulation of dipeptides, which correlated with the bacterial transition from a bacillary form to a coccoid morphology (23). Likewise, Möll *et al*. showed that the absence of the amidase AmiB, which is usually responsible for the cleavage of septal PG, led to an increase of dipeptides (~7% compared to < 1% for the WT) in *V. cholerae* (24). However, this deletion of *amiB* in *V. cholerae* resulted in filamentous cells due to their inability to divide properly (24), which is significantly distinguished from the ΔvarA phenotype described above. We therefore speculate that the altered PG in this mutant is not caused by a misregulated enzyme.

### WT-derived PG building blocks prevent rounding of ΔvarA cells

As we were unable to identify any obvious peptide-cleaving enzyme, we instead hypothesized that the altered PG composition of the ΔvarA cells could be caused by a lack of PG precursors. Irnov *et al*. demonstrated that changes in cell metabolism caused by a deletion of *hfq* in *Caulobacter crescentus* impaired the synthesis of meso-diaminopimelic acid (m-DAP). This in turn was shown to cause cell shape defects (25). Given that bacteria release PG building blocks into their surroundings during cell wall synthesis (26), we wondered whether the cell wall defect of the ΔvarA strain could be abrogated by WT-derived PG subunits. We therefore grew *varA*-deficient cells in WT-derived conditioned medium, which indeed prevented the rounding of the cells (Fig. 3A). To show that this phenotypic rescue was actually dependent on the recycling of the PG subunit, we removed the AmpG permease responsible for the import of muropeptides by generating a *varA/ampG*-deficient double mutant (27). As predicted, this double mutant could not be rescued by WT-derived conditioned medium (Fig. 3A). In contrast, a single *ampG*-deficient mutant showed no cell shape alterations compared to the WT (Fig. 3A).

**Figure 3.**
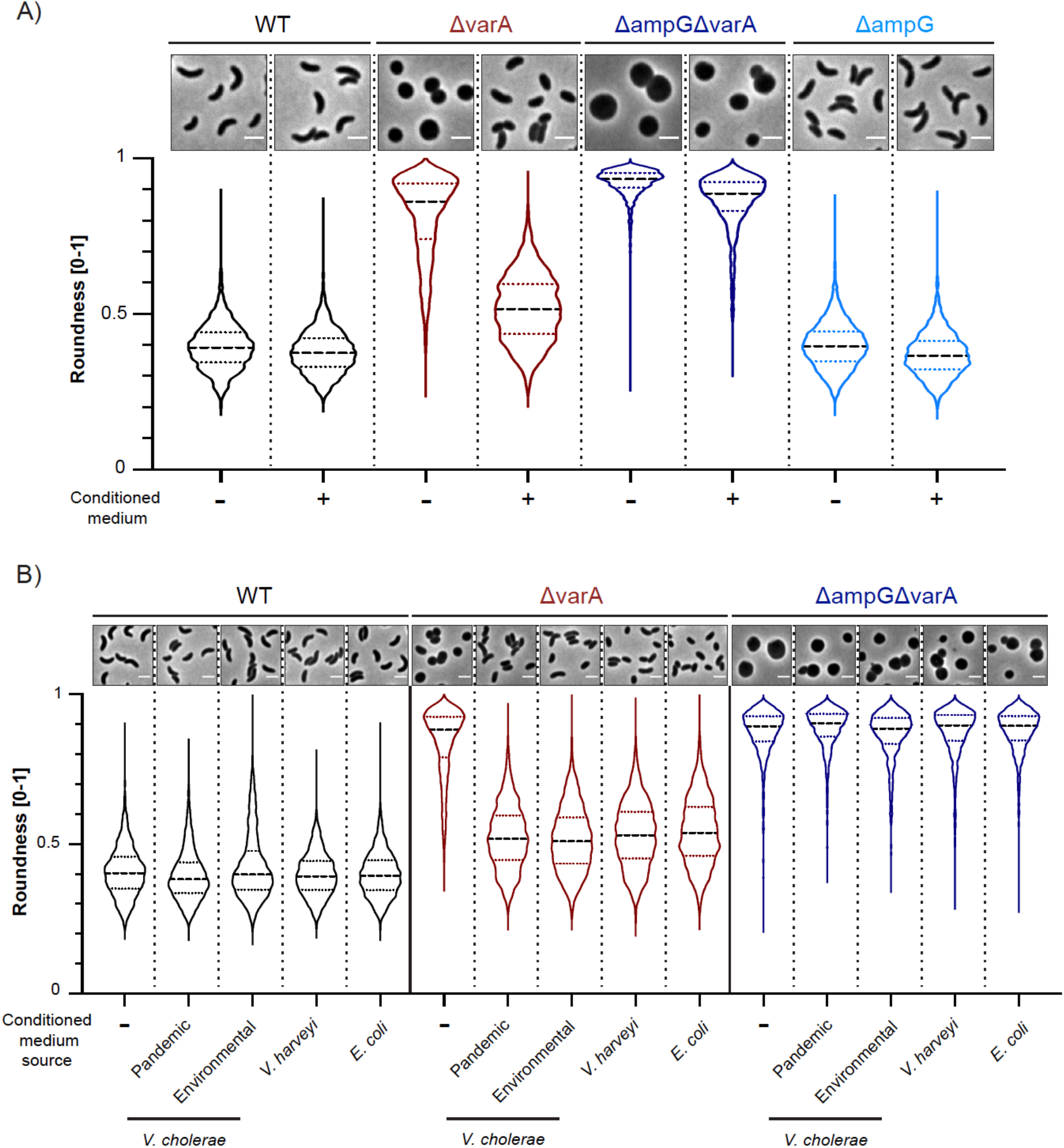
Recycling of PG prevents rounding of ΔvarA cells. (A) Conditioned medium rescues the *Vibrio* shape of ΔvarA in an AmpG-dependent manner. Phase contrast micrographs (top) and shape quantification (bottom) of the WT, ΔvarA, ΔampGΔvarA, and ΔampG strains grown in the absence (-) or presence (+) of conditioned medium. Scale bar: 2 μm. The quantification of roundness was performed with 3000 cells (n = 1000 per biologically independent experiment) using the MicrobeJ software. Roundness values range from 0 to 1. (B) Conditioned medium of diverse gram-negative bacteria rescues the ΔvarA rounding phenotype. Conditioned medium was derived from pandemic *V*. *cholerae* A1552 or environmental *V*. *cholerae* SA5Y, *V*. *harveyi*, or *E*. *coli*. Details as in panel A.

To study whether this phenotype was specific to the pandemic clade of *V. cholerae*, we used conditioned media derived from other bacteria, such as an environmental isolate of *V. cholerae* (strain SA5Y; (28, 29), *Vibrio harveyi* and *Escherichia coli*. As shown in Fig. 3B, conditioned media from any of these bacteria likewise prevented rounding of the ΔvarA cells. Collectively, these data support the notion that the shape transition of the ΔvarA strain is caused by a lack of cell wall precursors and that this phenotype can be rescued by PG recycling.

### Defects in *csrA* restore a normal cell shape for the ΔvarA strain

Based on the data provided above, we concluded that the cell rounding of *varA*-deficient cells was caused by a lack of cell wall precursors, which, ultimately, led to a dipeptide-enriched PG mesh with reduced cross-links. However, no link between VarA regulation and the synthesis of cell wall precursors had previously been described for *V. cholerae*. To better understand this connection and the underlying signaling pathway, we set up a transposon mutagenesis screen based on the assumption that the weakened cell wall of stationary phase ΔvarA cells would lyse under osmotic stress. Hence, we generated osmotic stress conditions by incubating the cells in NaCl-free LB medium (LB_0_), as previously reported (30). This significantly impacted the survival of the mutant cells, while WT cells remained unaffected (Fig. 4A).

**Figure 4.**
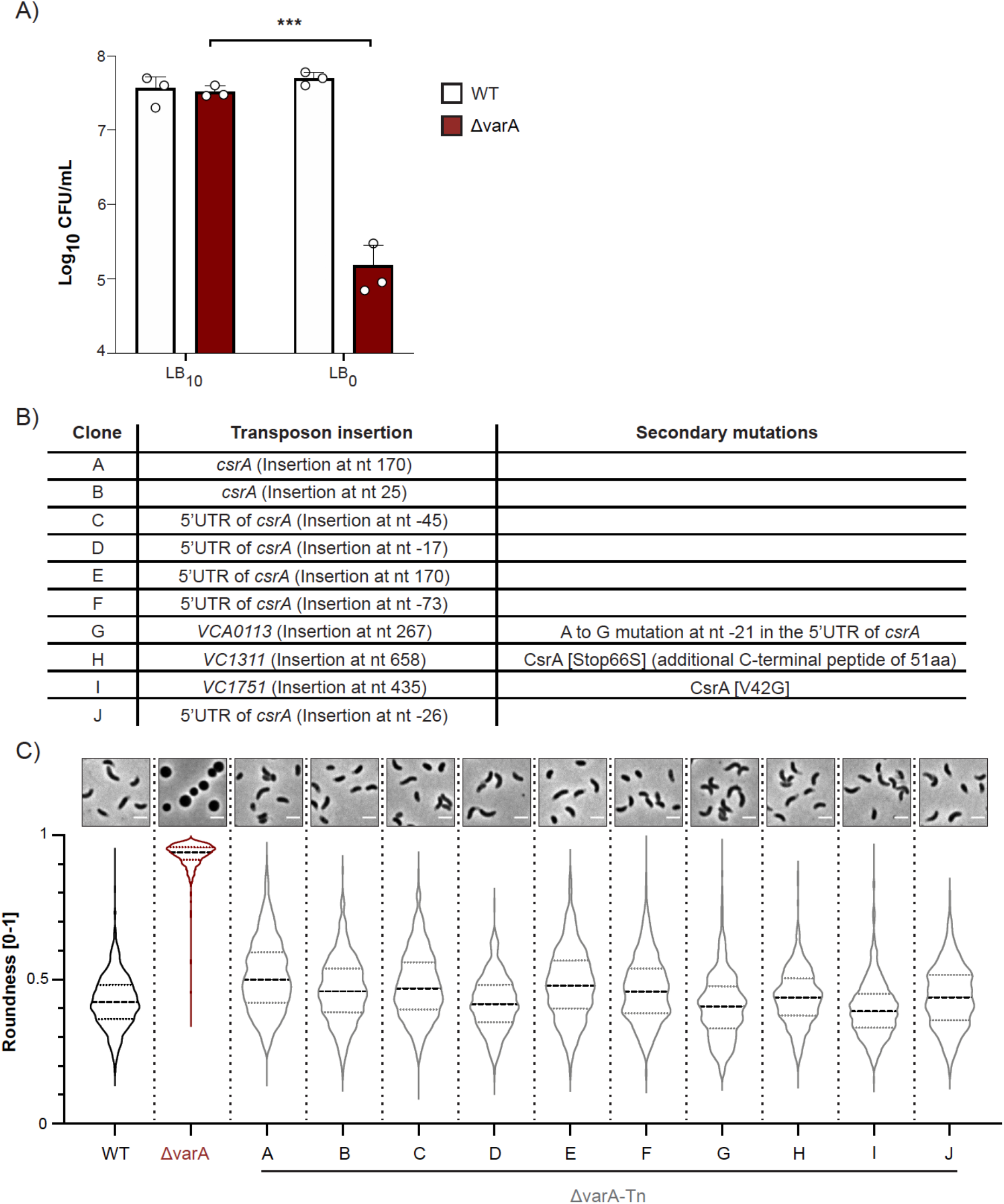
Suppressor mutants of ΔvarA are linked to *csrA*. (A) Late growth ΔvarA cells are sensitive to low osmolarity medium. After 20 h of growth, WT and ΔvarA bacteria were diluted 1:100 in regular LB_10_ or salt-free LB_0_ medium for 1 h before further dilution and plating to enumerate CFUs on the next day. Bars represent the mean of three independent repeats (error bars show the S.D.). (B) Summary of transposon hits. LB_0_-surviving mutants were isolated from the ΔvarA-Tn transposon library, and the transposon insertion sites were determined. Information for 10 different mutants is provided. (C) Suppressor mutants restored the *Vibrio* cell shape. Phase contrast micrographs (top) and roundness quantification (bottom) of the WT, ΔvarA, and the ten ΔvarA-Tn suppressor mutants (A-J). Cells were imaged at 20 h post-dilution. Imaging details as in Fig. 1. Scale bar: 2 μm.

To next identify suppressor mutants that would survive the LB_0_ treatment, we generated a mariner-based transposon library in the ΔvarA strain background; these mutants would have a normal PG, and therefore, a comma-shaped morphology. This library was subjected to three consecutive rounds of selection in LB_0_ medium to enrich surviving mutants. Following this experiment, we isolated 24 colonies and identified mutants containing 10 different transposon insertion sites (ΔvarA-Tn), as shown in Fig. 4B. Interestingly, 7 out of 10 mutants had the transposon inserted in or close to *csrA*. To check for any defects in *csrA* in the residual three mutants, we sequenced the *csrA* locus and its flanking regions. We discovered single nucleotide polymorphisms (SNPs) in all of them (Fig. 4B).

To confirm the selective advantage that these ten *csrA* suppressor mutants might have encountered during the exposure to osmotic stress, we assessed their cell morphology after overnight growth (Fig. 4C). Remarkably, all mutants exhibited a normal comma-like *Vibrio* morphology (Fig. 4C). As VarA inhibits CsrA function (4), we therefore concluded that the ΔvarA morphology was caused by increased CsrA activity due to a lack of VarA~P-dependent expression of the CsrA-sequestering sRNAs *csrB-D*. This finding is consistent with previous work showing that mutations in *csrA* suppressed other ΔvarA or ΔvarS phenotypes, including decreased glycogen storage in *V. cholerae* or growth defects in *V. cholerae* or a ΔgacA mutant of *Vibrio fischeri* (10, 11, 31). Collectively, these findings support the notion that the PG-dependent morphological changes are caused by increased CsrA activity in *varA*-deficient strains.

### Increased CsrA activity causes AspA overproduction

As CsrA is a post-transcriptional regulator (6), we wondered about potential changes at the protein level. We first looked at the global protein composition pattern using cell lysates of the WT and ΔvarA strains. As shown in Fig. 5A, a protein band with a size of around 50 kDa became starkly apparent for the *varA*-deficient sample after separation by SDS PAGE followed by Coomassie blue staining. Importantly, this band was diminished in the ten *csrA* suppressor mutants described above, supporting the role of CsrA in the overproduction of this protein (Fig. 5A).

**Figure 5.**
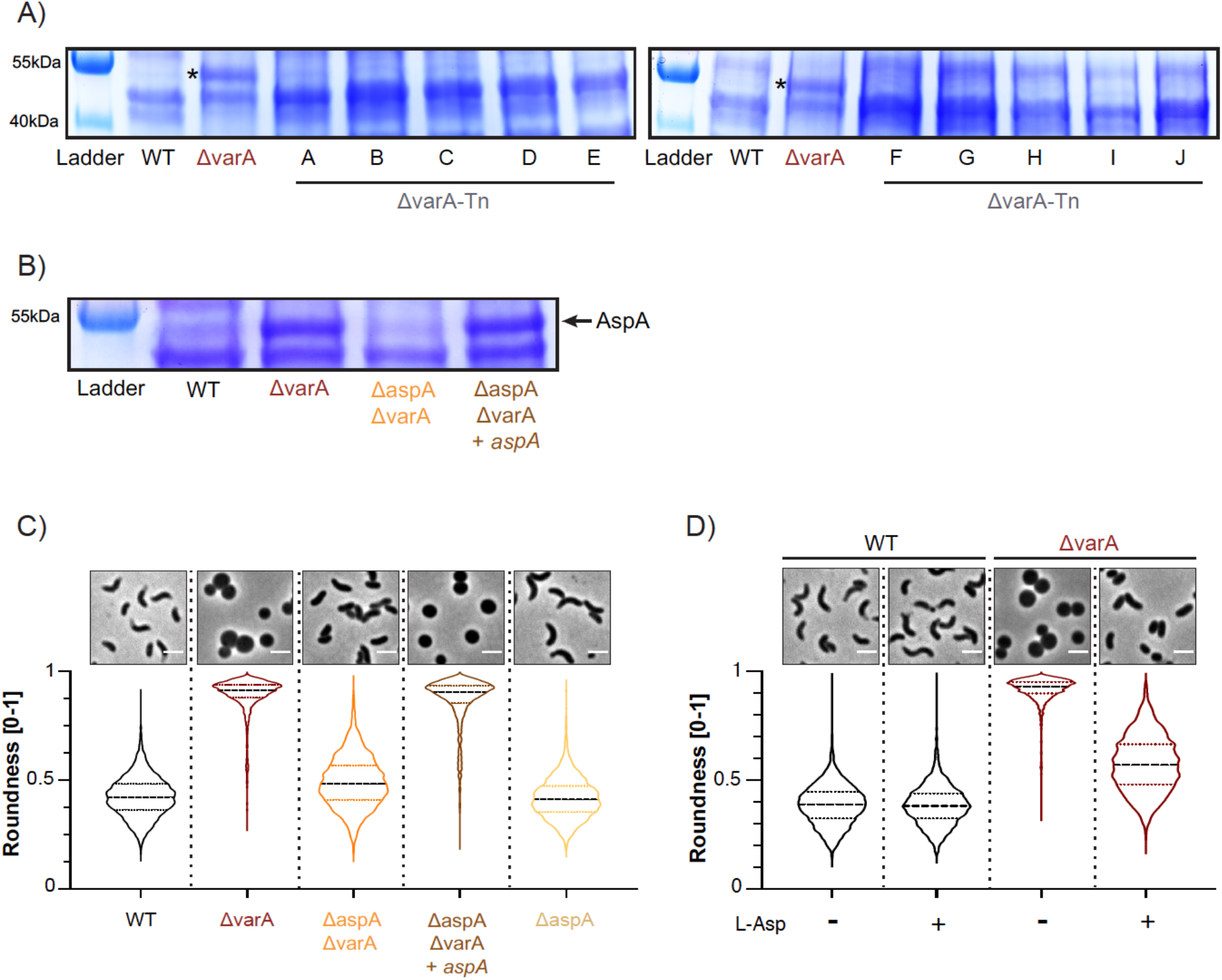
CsrA causes AspA overproduction and impairs m-DAP biosynthesis. (A) The protein pattern is different in the ΔvarA mutant. Coomassie blue staining of SDS PAGE-separated proteins from cell lysates of the WT, Δ*varA*, and the Δ*var*A-Tn suppressor mutants A-J. All strains were sampled at an OD_600_ of ~2.5. Bands of the protein ladder are shown in the first lane of each gel and their size is indicated on the left. (B) The dominant protein band corresponds to AspA. Coomassie blue staining of the total cell lysates of the WT, ΔvarA, ΔaspAΔvarA and the complemented strain ΔaspAΔvarA+*aspA*. Details as in panel A. The arrow points towards AspA. (C and D) Deletion of *aspA* or L-Asp supplementation abrogates rounding in ΔvarA cells. Phase contrast micrographs (top) and cell shape quantification (bottom) of the WT, ΔvarA, ΔaspAΔvarA, and ΔaspAΔvarA+*aspA* strains (C) or WT and ΔvarA cells grown in the absence (-) or presence (+) of L-aspartate (L-Asp). Cells were imaged at 20 h post-dilution. Imaging details as in Fig. 1 (n = 3000 cells for each condition). Scale bar: 2 μm.

Next, we analyzed this overproduced protein by mass spectrometry and identified it as aspartate ammonia lyase (AspA). Therefore, we deleted *aspA* in the ΔvarA background, resulting in the reversal of protein overproduction; additionally, this phenotype was able to be complemented by a new copy of *aspA* (ΔaspAΔvarA+*aspA*; Fig. 5B). Interestingly, previous studies identified the *aspA* mRNA amongst hundreds of direct CsrA targets in *Salmonella* using the CLIP-seq technique to identify protein-RNA interactions (32). We therefore conclude that CsrA enhances *aspA* mRNA translation in the *varA*-deficient *V. cholerae* mutant.

### AspA overproduction impairs the synthesis of m-DAP

To verify whether the overproduction of AspA was involved in the changed PG composition, therefore generating the round cell morphology of the ΔvarA mutant, we imaged the ΔaspAΔvarA strain after 20 h of growth. As shown in Fig. 5C, the *Vibrio* shape was restored in this double mutant, while the *aspA* complemented strain (ΔaspAΔvarA+*aspA*) retained the spherical morphology characteristic of the ΔvarA mutant. Deletion of *aspA* alone did not change the cellular morphology (Fig. 5C). Ergo, we conclude that AspA overproduction is involved in the PG-dependent rounding phenotype.

We next looked into how exactly AspA overproduction might affect morphology. AspA is responsible for the reversible conversion of aspartate into ammonia and fumarate (33). It is known that in many bacteria—including *V. cholerae*—L-aspartate is required for m-DAP synthesis as part of the lysine biosynthesis pathway (see KEGG online platform (34) accession number vch00300 for lysine biosynthesis in the *V. cholerae* reference strain N16961). Thus, we hypothesized that overproduction of AspA sequesters L-aspartate and, consequently, prevents m-DAP synthesis. To test this idea, we supplemented *varA-*deficient cells with L-aspartate. This significantly reduced cell rounding but did not alter the WT morphology (Fig. 5D), indicating that L-aspartate is indeed sequestered by AspA.

Taken all together, we propose a model whereby specific sequestering sRNAs are absent, causing an increased CsrA activity and the consequent overproduction of AspA, which reduces aspartate levels in ΔvarA cells (Fig. 6). These reduced aspartate levels subsequently impair the biosynthesis of the cell wall precursor m-DAP, thereby resulting in an abnormal dipeptide-containing under-crosslinked PG, which, ultimately, causes cell rounding (Fig. 6). Notably, until now, the signal(s) that abrogate(s) VarA phosphorylation by VarS, or any other additional histidine kinases, have not been unambiguously identified. Such conditions are expected to mimic the *varA*-deficient phenotypes that have been described in several previous studies (9–13), which we complement here with our finding of a VarA/CsrA-dependent PG modification and cell shape alteration. Thereupon, future studies are required to identify the conditions that alter VarA signaling during human infection and/or growth of *V. cholerae* in its natural aquatic habitat.

**Figure 6.**
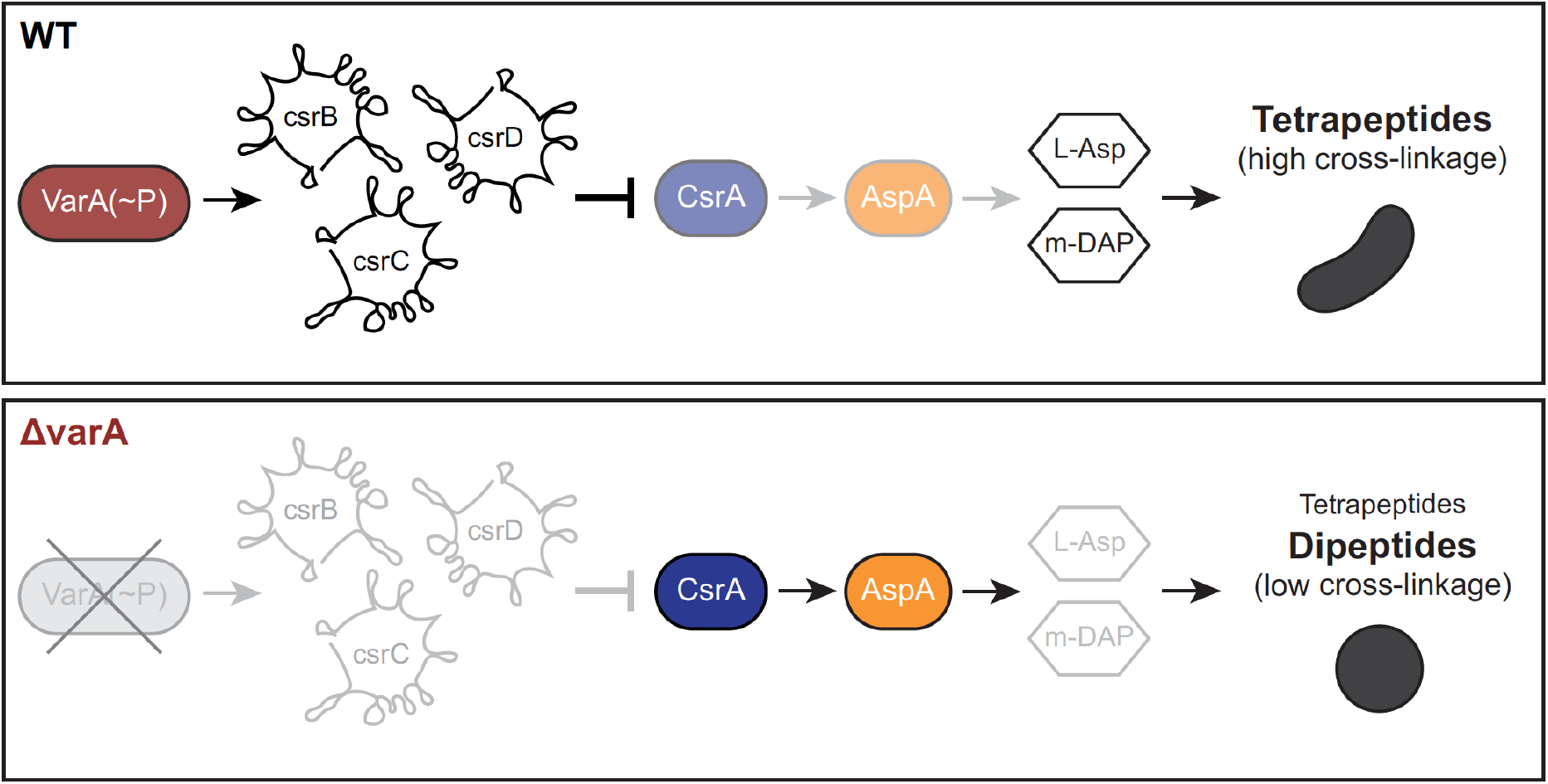
Model of VarA-CsrA signaling and impact on cell shape. For the WT, phosphorylated VarA [VarA(~P)] promotes the transcription of the sRNAs *csrB-D*, which sequesters CsrA. L-Asp and m-DAP levels are maintained resulting in normal PG and cell shape. In the absence of *varA*, the lack of the scavenging sRNAs results in increased CsrA activity and overproduced AspA, which reduces L-Asp levels and impairs m-DAP biosynthesis. The PG therefore contains dipeptides, while tetrapeptides are reduced, leading to a reduction of cross-links and therefore cell rounding.

Our study shows that impairment of the VarA-CsrA signaling pathway in *V. cholerae* leads to a modulation of bacterial morphology during the stationary growth phase by altering the synthesis of cell wall precursors. While the advantage of this alteration is so far unknown for *V. cholerae*, such cell shape transitions have been demonstrated by other pathogens, such as *C. jejuni* and *H. pylori* (22, 23). Moreover, growth phase-dependent cell wall remodeling occurs in many bacteria. For instance, during stationary phase, *V. cholerae* cells synthesize dedicated D-amino acids (D-aa), which are incorporated into their PG mesh (35). Production and insertion of such D-aa was hypothesized to be a strategy for adapting to changing conditions. Consistent with this idea, Le and colleagues recently demonstrated the incorporation of non-canonical D-aa into the PG of *Acinetobacter baumannii* during stationary phase (36). Interestingly, this PG editing process protected the pathogen from PG-targeting type VI secretion system-dependent effector proteins (36). Therefore, we postulate that round-shaped *V. cholerae* might encounter similar fitness benefits during interbacterial competition, antibiotic treatment, or other, so far unidentified, conditions.

Overall, our work has deciphered a new role of the VarA-CsrA signaling pathway in cell shape transition through the modulation of cell wall precursor synthesis. Due to the role of this pathway in bacterial virulence, understanding its purpose in cellular adaptation and survival is a key component of understanding bacterial changes that lead to human infections. The future studies we envision should therefore aim at identifying the conditions that lead to this altered signaling and conferred fitness benefits.

## Materials and Methods

### Bacterial strains and plasmids

The bacterial strains and plasmids used in this work are provided in Tables S2 and S3, respectively. The primary *V. cholerae* isolate used throughout this study is O1 El Tor strain A1552 (37), a representative of the ongoing 7^th^ cholera pandemic. Genetic manipulations are based on its published genome sequence (29).

### Growth conditions and medium supplementation

Bacterial cultures were grown aerobically with agitation (180 rpm) at 30 °C and 37 °C for *Vibrio* spp. and *E. coli*, respectively. As for the liquid medium, home-made lysogeny broth (10 g/L Tryptone, AppliChem; 10 g/L NaCl, Fisher scientific; 5 g/L Yeast Extract, AppliChem) was used or, when explicitly mentioned, a variation without NaCl (LB_0_). LB-agar (including 1.5% agar; Carl Roth) was used as a solid medium. All liquid cultures were grown overnight prior to being back-diluted (1:100) into fresh LB and were cultured for the indicated duration. When required, LB cultures were supplemented at 3h post-dilution with 7.5 mM L-aspartate (Sigma-Aldrich). Growth was monitored by measuring the optical density at 600 nm (OD_600_) and by enumerating the colony forming units (CFU) after serial dilution.

The conditioned medium was obtained from cultures grown for 8 h through centrifugation (4,000 rpm for 15 min at room temperature [RT]) and subsequent filter sterilization (0.2 μm filter) and were stored at 4 °C until the next day. Prior to usage, the conditioned medium was diluted 1:1 in two-fold concentrated LB medium (2× LB).

For natural transformation assays on chitin flakes, 0.5× defined artificial sea water (DASW) supplemented with 50 mM HEPES (Sigma-Aldrich) and vitamins (MEM, Gibco) was used, as described in (38). Thiosulfate citrate bile salts sucrose (TCBS; Sigma-Aldrich) agar plates were used to counter-select *E. coli* after bi-/or tri-parental mating. NaCl-free LB plates supplemented with 10% sucrose (Sigma-Aldrich) were used for the *sacB*-based counter-selection.

Whenever required, antibiotics were added at the following concentrations: ampicillin (Amp; 100 μg/ml), gentamicin (Gent; 50 μg/ml), kanamycin (Kan; 75 μg/ml), and rifampicin (Rif; 100 μg/ml).

### Genetic engineering of strains and plasmid constructions

Standard molecular biology-based techniques were used for molecular cloning (39). All genetically engineered strains were verified by PCR and confirmed by Sanger sequencing of the modified genomic regions (Microsynth AG, Switzerland). Deletion mutants of *V. cholerae* were generated via either an allelic exchange approach using the counter-selectable plasmid pGP704-Sac28 (40) or via a combination of natural transformation and Flip recombination to remove the selection cassette [TransFLP; (41–43)]. Tri-parental mating was used to site-directly integrate the mini-Tn7 transposon into the *V. cholerae* large chromosome (44). For complementation purposes, the transposon carried the gene-of-interest preceded by its promotor-containing upstream region.

### Light microscopy imaging

Thin agarose pads (1.2% in 0.5× PBS) were used to coat microscope slides. Bacteria were immobilized by mounting them on top of the pads, which were then covered with a coverslip. The cells were observed using a Zeiss Axio Imager M2 epi-fluorescence microscope, controlled by ZEN BLUE 2.6 software. All images were analyzed and prepared for publication using ImageJ (http://rsb.info.nih.gov/ij). For shape quantification purposes, single cell analysis was performed using MICROBEJ (15). For the latter approach, at least 10 images were taken per condition, from which 1000 bacterial cells were randomly selected for the final quantification. Each experiment was repeated three independent times.

### Cryo-electron microscopy

Restreaked colonies on LB-agar were used to inoculate LB medium to grow bacteria overnight. The resulting cultures were then back-diluted (1:100) into fresh LB medium. At 20 h post-dilution, an aliquot was removed, mixed with 15 nm gold beads (Cell Microscopy Core, Utrecht University, Utrecht, The Netherlands), loaded onto a R2/2 200 mesh Cu grid (Quantifoil Micro Tools, GmbH), and plunge frozen in liquid ethane using the automated Leica EM GP plunge freezer (Leica Microsystems GmbH). Vitrified cells were imaged using a Gatan 626 cryoholder using a Talos L120C electron microscope (Thermo Fisher Scientific, TFS). Images were acquired with a Ceta CMOS camera (TFS) with a magnification range of 1,600– 17,500×, corresponding to a pixel size of 6.29–0.584 nm/pixel (FOV 25.16–2.336 μm^2^).

### Peptidoglycan analysis

PG was isolated from two biological replicates and analyzed by High Performance Liquid Chromatography (HPLC), as previously described (45). Cells were grown at 30 °C in 400 mL of LB medium with agitation for 2 h and 20 h. Cells were cooled on ice for 15 min and were collected by centrifugation for 15 min at 4 °C and 3,220× *g*. Cell pellets were resuspended in 6 mL of cold water and lysed by adding them dropwise to 6 mL of boiling 8% SDS (Sigma-Aldrich) within 10 min, under vigorous stirring. Samples were boiled for an additional 30 min and subsequently cooled to RT and stored until further analysis.

Crude PG was collected by centrifugation for 60 min at 90,000× *g* at 28°C. Pellets were washed several times with warm water until the SDS was removed. Samples were treated with α-amylase and pronase to remove high–molecular weight glycogen and PG-associated proteins, respectively. The resulting PG was boiled in 4% SDS and was washed free of SDS as described (45).

The PG composition (muropeptide profile) was analyzed as previously described (45). Briefly, muropeptides were generated from PG using the muramidase cellosyl (Hoechst, Frankfurt am Main, Germany). The reaction was stopped by heating the samples at 100 °C for 10 min and the sample was centrifuged for 10 min at 13,000× *g* to clarify the solution. Muropeptides present in the supernatant were reduced with sodium borohydride and analyzed by HPLC using a 250 × 4.6 mm, 3 μm ProntoSIL 120-3-6C18 AQ reversed phase column (Bischoff, Leonberg, Germany) on an Agilent 1100 system. The eluted muropeptides were detected by their absorbance at 205 nm. *V. cholerae* muropeptides were assigned according to their known retention times and quantified by their peak area using the Laura software (Lab Logic Systems). The muropeptide fraction 4 was collected and analyzed by mass spectrometry at the Newcastle University Pinnacle facility as described previously (46).

### Transposon mutagenesis screen

Transposon insertion libraries were prepared in the ΔvarA strain by introducing the mariner-based transposon (Kan^R^) carried on plasmid pSC189 (47) via conjugation, as previously described (48). Following growth at 30°C overnight, colonies (~100,000) were scrapped from the plates and resuspended in PBS buffer. To screen for mutants resistant to osmotic stress, the library was diluted 1:100 in LB_0_ plus kanamycin and was incubated at 30 °C for 1 h with agitation. Surviving bacteria were concentrated by centrifugation (3,220× *g* for 15 min at RT), resuspended in regular LB_10_ medium (containing kanamycin) and cultured at 30°C overnight. Following two additional rounds of selection in LB_0_ medium, the library was stored in LB medium containing 20% glycerol at −80 °C until further analysis.

Next, the libraries were thawed, mixed with LB_10_ + Kan, and grown at 30 °C overnight before being back-diluted 1:100 in fresh medium and grown for approximately 20 h. After visualization of the bacterial cell morphology by microscopy to confirm the absence of round cells, the resulting cultures were streaked on LB-agar plates (containing kanamycin) to obtain single colonies. To identify transposon insertion sites, 24 colonies were randomly picked and subjected to two-step arbitrary PCR, followed by Sanger sequencing to identify the transposition site, as previously described (48).

### SDS-PAGE, Western blotting and Coomassie blue staining

Bacteria were grown as described above until they reached an OD_600_ of approximately 2.5. At that point, the cultures were harvested by centrifugation (20,000 × *g* for 3 min at RT) and the pellet was resuspended in a volume of 2× Laemmli buffer (Sigma-Aldrich) that adjusts for the total number of bacteria according to the OD_600_ of the initial cultures followed by 15 min incubation at 95 °C. The proteins were resolved by SDS-PAGE using 8–16% Mini-PROTEAN TGX Stain-Free protein gels (Bio-Rad) and transferred onto a PVDF membrane using a semi-dry apparatus (Trans-Blot Turbo Transfer System; Bio-Rad). The detection of the signal from the HapR protein or Sigma70 as a loading control was performed as described (48).

Coomassie blue staining was performed after SDS-PAGE to identify under- or overproduced proteins. To do so, the gels were soaked in the Coomassie blue solution (0.2% Coomassie brilliant blue [A1092; AppliChem] in 10% methanol plus 1% acetic acid) and stained for 30 min at RT with gentle agitation. Three destaining steps (30 min each) were performed using a destaining solution (10% methanol plus 1% acetic acid).

### Protein identification through mass spectrometry

Gel pieces previously stained with Coomassie blue as described above were washed twice in 50% ethanol and 50 mM ammonium bicarbonate (Sigma-Aldrich) for 20 min and dried by vacuum centrifugation. Sample reduction was performed with 10 mM dithioerythritol (Merck-Millipore) for 1 h at 56 °C. This washing-drying step was repeated before performing the alkylation step with 55 mM iodoacetamide (Sigma-Aldrich) for 45 min at 37°C in the dark. Next, samples were washed-dried once again and digested overnight at 37°C using mass spectrometry grade Trypsin gold (Trypsin Gold, Promega) at a concentration of 12.5 ng/μl in 50 mM ammonium bicarbonate and 10 mM CaCl_2_. The resulting peptides were extracted in 70% ethanol plus 5% formic acid (Merck-Millipore) twice for 20 min with permanent shaking. Samples were further dried by vacuum centrifugation and stored at −20 °C. Peptides were desalted on C18 StageTips (49) and dried by vacuum centrifugation prior to LC-MS/MS injections. Samples were resuspended in 2% acetonitrile (Biosolve) and 0.1% formic acid, and nano-flow separations were performed on a Dionex Ultimate 3000 RSLC nano UPLC system (Thermo Fischer Scientific) online connected with a Q Exactive HF Orbitrap mass spectrometer (Thermo Fischer Scientific). A capillary precolumn (Acclaim Pepmap C18, 3 μm-100 Å, 2 cm × 75 μm ID) was used for sample trapping and cleaning. A 50 cm long capillary column (75 μm ID; in-house packed using ReproSil-Pur C18-AQ 1.9 μm silica beads; Dr. Maisch) was then used for analytical separations at 250 nl/min over 90 min biphasic gradients. Acquisitions were performed through Data-Dependent Acquisition (DDA). First MS scans were acquired with a resolution of 60,000 (at 200 m/z) and the 15 most intense parent ions were then selected and fragmented by High energy Collision Dissociation (HCD) with a Normalized Collision Energy (NCE) of 27% using an isolation window of 1.4m/z. Fragmented ions were acquired with a resolution of 15,000 (at 200m/z) and selected ions were then excluded for the following 20 s. Raw data were processed using SEQUEST in Proteome Discoverer v.2.2 against a home-made database (3610 entries) using the genome sequence of *V. cholerae* strain A1552 as an input (GenBank accession numbers CP028894 for chromosome 1 and CP028895 for chromosome 2) (29). Enzyme specificity was set to trypsin and a minimum of six amino acids were required for peptide identification. Up to two missed cleavages were allowed. A 1% FDR cut-off was applied both at peptide and protein identification levels. For the database search, carbamidomethylation was set as a fixed modification, whereas oxidation (M), acetylation (protein N-term), PyroGlu (N-term Q), and Phosphorylation (S,T,Y) were considered as variable modifications. Data were further processed and inspected in Scaffold 4.10 (Proteome Software, Portland, USA).

## Acknowledgements

We thank Daniela Vollmer for purification of PG, Joe Gray for muropeptide analysis by mass spectrometry, members of the proteomics core facility within the School of Life Sciences at EPFL for the AspA identification, and John Mekalanos & Andrew Camilli for provision of *V. cholerae* strains. We also acknowledge Nicolas Flaugnatti for help with the preparation of the SDS-boiled cell lysates, and Milena Jaskólska & David W. Adams for their contribution to the mentoring of L.F.L.R.. We thank Justine Collier and Tobias Dörr for advice and members of the Blokesch laboratory for valuable discussions. This work was supported by EPFL intramural funding and a Consolidator Grant from the European Research Council (ERC; 724630-CholeraIndex) to M.B. and by a Wellcome Trust Senior Investigator Award (101824/Z/13/Z) to W.V.. M.B. is a Howard Hughes Medical Institute (HHMI) International Research Scholar (#55008726).

