## Supporting Information for "The VarA-CsrA regulatory pathway influences cell shape in *Vibrio cholerae*"

**This PDF file includes:**

Supporting Figures S1 to S4

Supporting Tables S1 to S3

Supporting References

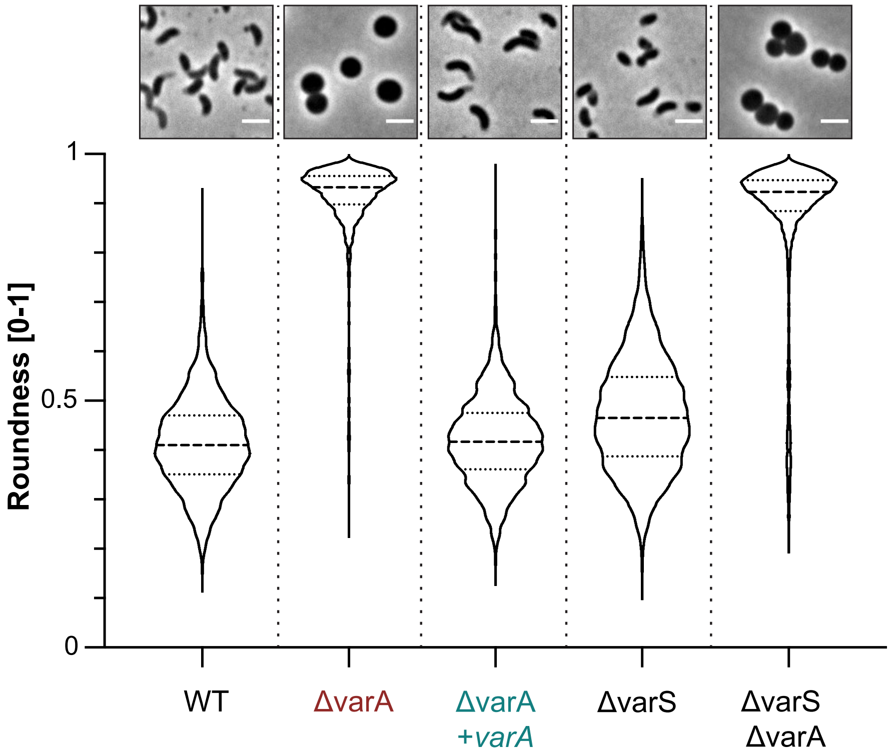


**Figure S1. *V. cholerae* strains lacking VarA but not VarS display an atypical round shape late during growth.** Phase contrast micrographs (top) and roundness quantification (bottom) of the WT, ΔvarA, ΔvarS, complemented ΔvarA+*varA*, and ΔvarSΔvarA strains. Cells were imaged at 20 h post-dilution. Scale bar: 2 µm. The roundness quantification is based on 3000 cells (n = 1000 per independent repeat) using the MicrobeJ software.

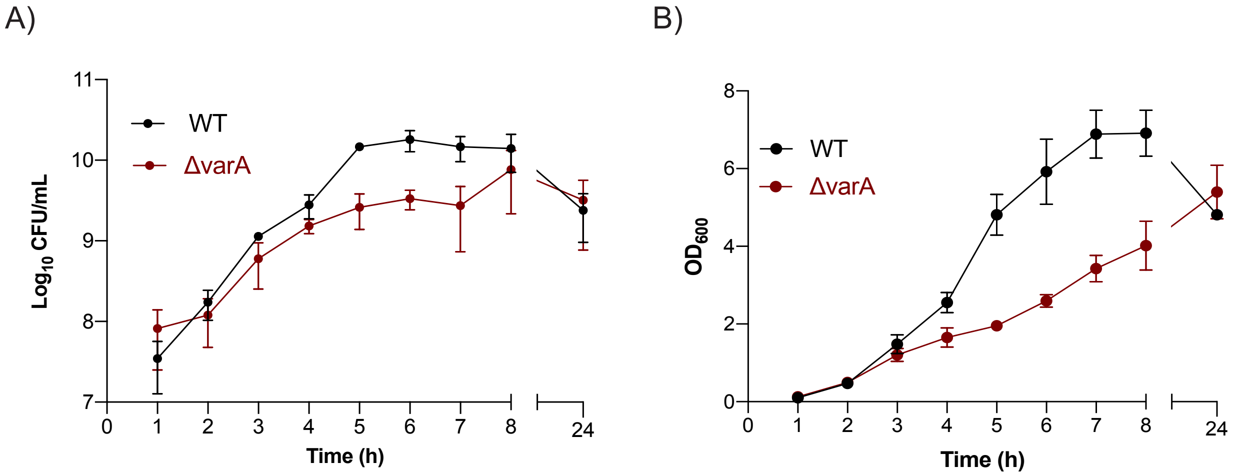


**Figure S2. The ΔvarA mutant has a slight growth defect.** Growth curves of the WT and ΔvarA strains. (A) The colony forming units (CFU) (B) and the optical density at 600 nm ($OD_{600}$) were measured every hour for 8 h and again at 24 h post-dilution. Each value represents the mean of 3 independent experiments (± S.D., as shown by the error bars).

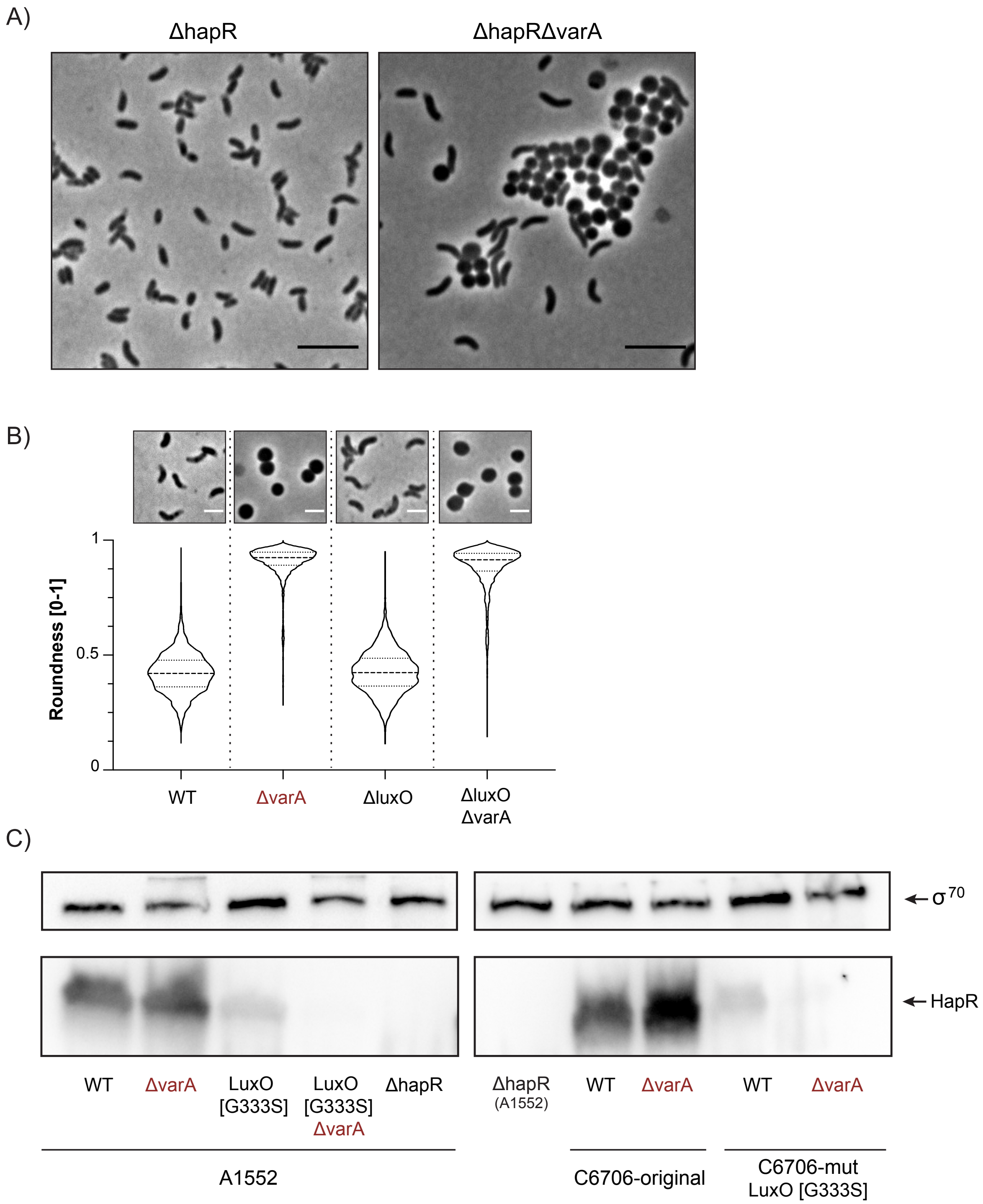


**Figure S3. The ΔvarA mutant is QS-proficient.** (A and B) HapR is produced and active in ΔvarA strains. Phase contrast micrographs of the following strains after growth for 20 h: ΔhapR and ΔhapRΔvarA strains (A; scale bar: 5 µm) or WT, ΔvarA, ΔluxO, and ΔluxOΔvarA (B; scale bar: 2 µm). Roundness quantification of n = 3000 cells for each condition is provided in (B). (C) QS-impaired LuxO* variants abrogate HapR production. Detection of HapR by western blotting for WT and *luxO** variants of strains A1552 and C6706 in the presence or absence of *varA*. All strains were sampled at $OD_{600}$ ~ 2.5. Details as in Fig. 1C.

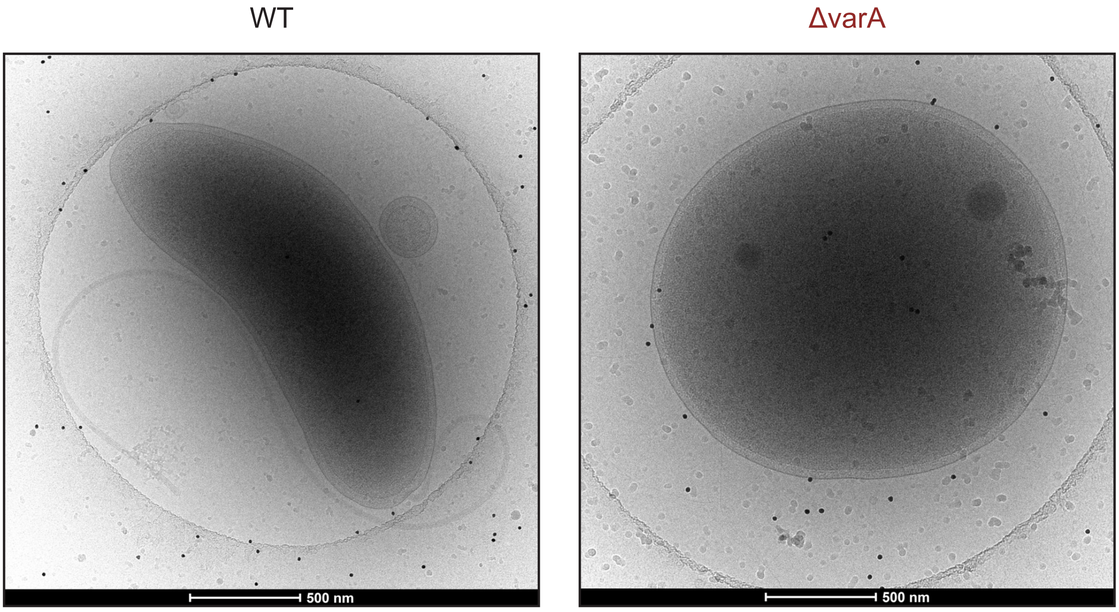


**Figure S4. ΔvarA cells do not show periplasmic or membrane defects.** Ultrastructural analysis of WT and ΔvarA cells imaged by cryo-electron microscopy. The cells were sampled at 20 h post-dilution. Scale bar: 500 nm.

**Table S1: Summary of the muropeptide composition of diverse *V. cholerae* strains.**

| Muropeptide[2] | Relative % of muropeptide[1] | | | | | |
|---|---|---|---|---|---|---|
| | 2 h post-dilution | | | 20 h post-dilution | | |
| | WT | Δ*varA* | ΔvarA +*varA* | WT | Δ*varA* | ΔvarA +*varA* |
| **Tri** | 0.7 ± 0.2 | 1.6 ± 0.9 | 0.6 ± 0.0 | 2.0 ± 0.3 | 4.8 ± 0.4 | 3.0 ± 0.3 |
| **TetraGly4** | 1.2 ± 0.4 | 2.3 ± 1.1 | 0.8 ± 0.1 | 1.0 ± 0.2 | 1.2 ± 0.1 | 1.5 ± 0.0 |
| **Tetra** | 49.4 ± 1.0 | 46.1 ± 3.1 | 49.4 ± 0.8 | 42.6 ± 2.9 | 27.7 ± 1.1 | 40.4 ± 0.5 |
| **Di[3]** | 0.0 ± 0.0 | 1.7 ± 0.4 | 0.0 ± 0.0 | 0.0 ± 0.0 | 24.4 ± 0.0 | 0.0 ± 0.0 |
| **Penta** | 2.8 ± 0.4 | 2.2 ± 0.7 | 3.1 ± 0.5 | 0.6 ± 0.1 | 2.5 ± 0.4 | 0.8 ± 0.3 |
| ***unknown*** | 0.7 ± 0.1 | 1.4 ± 0.2 | 0.8 ± 0.1 | 2.0 ± 1.1 | 4.5 ± 0.9 | 2.4 ± 0.5 |
| **TriTetra** | 0.6 ± 0.2 | 3.5 ± 2 | 0.4 ± 0.4 | 1.6 ± 0.3 | 3.4 ± 1.2 | 4.4 ± 2.6 |
| **TetraTri** | 0.3 ± 0.3 | 1.1 ± 0.2 | 0.3 ± 0.3 | 1.6 ± 0.3 | 2.5 ± 0.2 | 2.1 ± 0.2 |
| **TetraTetra** | 30.2 ± 0.0 | 27.4 ± 0.6 | 30.5 ± 0.5 | 29.0 ± 0.3 | 14.8 ± 0.6 | 27.8 ± 0.6 |
| **TetraPenta** | 1.0 ± 0.2 | 0.8 ± 0.0 | 1.2 ± 0.2 | 1.0 ± 0.5 | 1.3 ± 0.2 | 1.0 ± 0.0 |
| **TetraTetraTetra** | 1.6 ± 0.2 | 1.5 ± 0.4 | 1.4 ± 0.1 | 1.7 ± 0.1 | 1.3 ± 0.1 | 1.5 ± 0.1 |
| **TetraTetraAnh I** | 6.8 ± 0.9 | 5.2 ± 1.7 | 6.6 ± 0.4 | 6.5 ± 1.9 | 4.7 ± 2.0 | 5.6 ± 2.0 |
| **TetraTetraAnh II** | 3.2 ± 0.2 | 3.9 ± 1.6 | 3.4 ± 0.5 | 7.3 ± 0.4 | 4.7 ± 0.5 | 6.9 ± 0.3 |
| **TetraTetraTetraAnh I** | 1.7 ± 0.0 | 1.4 ± 0.4 | 1.9 ± 0.3 | 3.1 ± 0.3 | 2.2 ± 0.5 | 2.6 ± 0.4 |
| ***Summary*** | | | | | | |
| **Monomers** | 54.7 ± 1.1 | 55.2 ± 2.3 | 54.6 ± 0.4 | 48.1 ± 1.7 | 65.1 ± 1.1 | 48.1 ± 0.5 |
| **Dimers** | 42.0 ± 0.9 | 41.9 ± 1.6 | 42.4 ± 0.1 | 47.0 ± 1.3 | 31.4 ± 0.8 | 47.8 ± 0.1 |
| **Trimers** | 3.3 ± 0.2 | 2.9 ± 0.7 | 3.1 ± 0.4 | 4.9 ± 0.4 | 3.5 ± 0.2 | 4.1 ± 0.6 |
| **Dipeptide (total)** | 0.0 ± 0.0 | 1.7 ± 0.4 | 0.0 ± 0.0 | 0.0 ± 0.0 | 24.4 ± 0.0 | 0.0 ± 0.0 |
| **Tripeptide (total)** | 3.0 ± 0.8 | 7.6 ± 1.2 | 2.5 ± 0.6 | 6.6 ± 0.8 | 13.5 ± 0.7 | 10.1 ± 0.4 |
| **Tetrapeptide (total)** | 93.7 ± 2.5 | 88.2 ± 1.2 | 93.9 ± 0.3 | 92.4 ± 5.9 | 59.1 ± 3.9 | 88.6 ± 2.7 |
| **Pentapeptide (total)** | 3.3 ± 0.5 | 2.6 ± 0.7 | 3.7 ± 0.6 | 1.1 ± 0.2 | 3.1 ± 0.5 | 1.3 ± 0.3 |
| **Peptides in cross-linkage (%)** | 45.3 ± 1.1 | 44.8 ± 2.3 | 45.4 ± 0.4 | 51.9 ± 1.7 | 34.9 ± 1.1 | 51.9 ± 0.6 |
| **Average chain length (DS)** | 18.1 ± 1.3 | 19.9 ± 0.2 | 18.1 ± 0.4 | 12.6 ± 1.5 | 18.4 ± 3.1 | 14.0 ± 2.3 |

[1] Values are means ± variation for two independent PG preparations. The relative peak areas were estimated as the percentage of all known peaks.

[2] Nomenclature of muropeptides as in (1).

[3] The Di fraction of sample Δ*varA*-20 h was collected and confirmed by mass spectrometry. The measured neutral mass was 698.0079 amu, the theoretical mass of GlcNAc-MurNAc(red)-L-Ala-D-Glu (Di) is 698.2558 amu.

### Table S2. Bacterial strains used in this study

| Strains | Genotype and description | Internal strain number | Reference |
|---|---|---|---|
| ***Vibrio cholerae*** | | | |
| WT | Wild-type A1552 O1 El Tor Inaba; Rif$^R$ | GC#1 | (2) |
| ΔvarA | A1552 deleted for *varA* (*VC1213*) via TransFLP; Rif$^R$ | GC#3812 | This study |
| ΔvarA+*varA* | A1552Δ*varA* containing mini-Tn7-*varA*; Rif$^R$, Gent$^R$ | GC#9263 | This study |
| ΔvarS | A1552 deleted for *varS* (*VC2453*) via TransFLP; Rif$^R$ | GC#3814 | This study |
| ΔvarSΔvarA | A1552Δ*varS* deleted for *varA* (*VC1213*) using suicide plasmid pGP704-Sac28-Δ*varA*; Rif$^R$ | GC#9264 | This study |
| ΔhapR | A1552Δ*hapR* (*VC0583*) | GC#3 | (3) |
| ΔhapRΔvarA | A1552Δ*hapR* deleted for *varA* (*VC1213*) using suicide plasmid pGP704-Sac28-Δ*varA*; Rif$^R$ | GC#9265 | This study |
| ΔluxO | A1552Δ*luxO* (*VC1021*); Rif$^R$ | GC#20 | (3) |
| ΔluxOΔvarA | A1552Δ*luxO* deleted for *varA* (*VC1213*) via TransFLP; Rif$^R$ | GC#9267 | This study |
| ΔampG | A1552 deleted for *ampG* (*VC2300*) via TransFLP; Rif$^R$ | GC#9275 | This study |
| ΔampGΔvarA | A1552Δ*ampG* deleted for *varA* (*VC1213*) via TransFLP; Rif$^R$ | GC#9277 | This study |
| ΔaspA | A1552 deleted for *aspA* (*VC2698*) via TransFLP; Rif$^R$ | GC#9269 | This study |
| ΔaspAΔvarA | A1552ΔaspA::FRT deleted for *varA* (*VC1213*) via TransFLP; Rif$^R$ | GC#9272 | This study |
| ΔaspAΔvarA+*aspA* | A1552ΔaspAΔ*varA* containing mini-Tn7-*aspA*; Rif$^R$, Gent$^R$ | GC#9273 | This study |
| luxO[G333S] | A1552 with site-directed point mutation in *luxO* (resulting in LuxO[G333S]) | GC#4492 | (4) |
| luxO[G333S]ΔvarA | A1552-*luxO*[G333S] deleted for *varA* (*VC1213*) via TransFLP; Rif$^R$ | GC#9284 | This study |
| C6706-original | Wild-type C6706; O1 El Tor Inaba; non-mutated *luxO*; Str$^S$ | GC#4522 | Gift from J. Mekalanos; (4); |
| C6706Δ*varA* | C6706 deleted for *varA* (*VC1213*) via TransFLP; Str$^S$ | GC#9280 | This study |
| C6706 (here: C6706-mut) | C6706 lacZ+ strain with mutated *luxO*[G333S]; Str$^R$ | GC#4524 | Gift from J. Mekalanos (before repair; see (5)) |
| C6706-mutΔvarA | C6706-mut deleted for *varA* (*VC1213*) via TransFLP; Str$^R$ | GC#9282 | This study |
| E7946 | Wild-type E7946; O1 El Tor Ogawa; isolated in 1978, Bahrain; Str$^R$ | GC#2600 | Gift from A. Camilli (6) |
| E7946ΔvarA | E7946 deleted for *varA* (*VC1213*) via TransFLP; Str$^S$ | GC#9278 | This study |
| E7946 (here: E7946-AC) | Wild-type E7946; O1 El Tor Ogawa; isolated in 1978, Bahrain; Str$^R$ | GC#6824 | Gift from A. Camilli (6) |
| E7946Δ*varA* (here: E7946-AC Δ*varA*) | E7946 deleted for *varA* (*VC1213*); Str$^R$ | GC#6825 | Gift from A. Camilli; (7) |

| | | | |
|---|---|---|---|
| SA5Y | Environmental *V. cholerae* isolate collected in Old Salinas River (CA, USA) in May 2004 | GC#353 | (8) |
| ***V. cholerae*** **transposon mutants** | | | |
| ΔvarA-Tn clone A | A1552ΔvarA carrying the *mariner*-based transposon inserted in *csrA* | GC#9285 | This study |
| ΔvarA-Tn clone B | A1552ΔvarA carrying the *mariner*-based transposon inserted in *csrA* | GC#9286 | This study |
| ΔvarA-Tn clone C | A1552ΔvarA carrying the *mariner*-based transposon inserted in the upstream region of *csrA* | GC#9287 | This study |
| ΔvarA-Tn clone D | A1552ΔvarA carrying the *mariner*-based transposon inserted in the upstream region of *csrA* | GC#9288 | This study |
| ΔvarA-Tn clone E | A1552ΔvarA carrying the *mariner*-based transposon inserted in the upstream region of *csrA* | GC#9289 | This study |
| ΔvarA-Tn clone F | A1552ΔvarA carrying the *mariner*-based transposon inserted in the upstream region of *csrA* | GC#9290 | This study |
| ΔvarA-Tn clone G | A1552ΔvarA carrying the *mariner*-based transposon inserted in *VCA0113;* A to G mutation at position -21 of *csrA* | GC#9291 | This study |
| ΔvarA-Tn clone H | A1552ΔvarA carrying the *mariner*-based transposon inserted in *VC1311;* A197C mutation in *csrA* resulting in CsrA[Stop66S] | GC#9292 | This study |
| ΔvarA-Tn clone I | A1552ΔvarA carrying the *mariner*-based transposon inserted in *VC1751;* T124C mutation in *csrA* resulting in CsrA[V42G] | GC#9293 | This study |
| ΔvarA-Tn clone J | A1552ΔvarA carrying the *mariner*-based transposon inserted in the upstream region of *csrA* | GC#9294 | This study |
| ***Vibrio harveyi*** | | | |
| *Vibrio harveyi* DSM-19623 | *Vibrio harveyi* WT strain from DSMZ | GC#3440 | DSMZ culture collection |
| ***Escherichia coli*** | | | |
| S17-1λpir | $Tp^R$ $Sm^R$ *recA thi pro hsdR-M+ RP4:2-Tc:Mu*: $Km^R$ Tn7 (λ*pir*) | GC#648 | (9) |
| MG1655 | F- lambda- *ilvG- rfb-50 rph-1* (cured derivative of K-12) | GC#2904 | Laboratory collection |
| MFD*pir* | MG1655 RP4-2-Tc::[Mu1::*aac*(3)IV-Δ*aphA*- Δ*nic35*-ΔMu2::zeo] Δ*dapA*::(erm-pir) Δ*recA* | GC#4662 | (10) |

**Table S3: Plasmids used in this study.**

| Plasmid | Genotype | Internal strain number | Reference |
|---|---|---|---|
| pBR-FRT-Kan-FRT2 | pBR322 derivative containing improved FRT-*aph*-FRT cassette, used as template for TransFLP; $Amp^R$, $Kan^R$ | GC#3782 | (11) |
| pBR-flp | pBR322 derivative containing FLP+, λ cI857+, λ pR from pCP20 integrated into the *EcoRV* site of pBR322; used for FLP recombination; $Amp^R$ | GC#1203 | (12) |
| pUX-BF13 | pUX-BF13-oriR6K, helper plasmid with Tn7 transposition function; $Amp^R$ | GC#457 | (13) |
| pGP704-mTn7 | pGP704 with mini-Tn7; $Amp^R$, $Gent^R$ | GC#645 | (14) |
| pGP704-Sac28 | suicide vector, *ori R6K*, *sacB*; $Amp^R$ | GC#649 | (15) |
| pSC189 | Plasmid for the delivery of *mariner*-based transposon; $Amp^R$, $Kan^R$ | GC#6089 | (16) |
| pGP704-Sac28ΔvarA | pGP704-Sac28 with *varA* gene fragment (ΔvarA::FRT ; derived from the *varA* deletion via TransFLP) and its flanking regions; $Amp^R$ | GC#9261 | This study |
| pGP704-mTn7-*varA* | pGP704 with mini-Tn7 carrying *varA* and its 164bp upstream region (with potential promoter region); $Amp^R$ | GC#9260 | This study |
| pGP704-mTn7-*aspA* | pGP704 with mini-Tn7 carrying *aspA* and its 247bp upstream region (with potential promoter region); $Amp^R$ | GC#9262 | This study |
